# Switch-like and persistent memory formation in individual *Drosophila* larvae

**DOI:** 10.1101/2021.04.15.440041

**Authors:** Amanda Lesar, Javan Tahir, Jason Wolk, Marc Gershow

## Abstract

Associative learning allows animals to use past experience to predict future events. The circuits underlying memory formation support immediate and sustained changes in function, often in response to a single example. Larval *Drosophila* is a genetic model for memory formation that can be accessed at molecular, synaptic, cellular, and circuit levels, often simultaneously, but existing behavioral assays for larval learning and memory do not address individual animals, and it has been difficult to form long lasting memories, especially those requiring synaptic re-organization. We demonstrate a new assay for learning and memory capable of tracking the changing preferences of individual larvae. We use this assay to explore how activation of a pair of reward neurons changes the response to the innately aversive gas Carbon Dioxide (CO_2_). We confirm that when coupled to odor presentation in appropriate temporal sequence, optogenetic reward reduces avoidance of CO_2_. We find that learning is switch-like: all-or-none and quantized in two states. Memories can be extinguished by repeated unrewarded exposure to CO_2_ but are stabilized against extinction by repeated training or overnight consolidation. Finally, we demonstrate long-lasting protein synthesis dependent and independent memory formation.

## Introduction

Associative learning allows animals to use past experience to predict important future events, such as the appearance of food or predators, or changes in their environmental conditions [1, 2]. The *Drosophila* larva is a favorable model system for the study of learning and memory formation [3–9], with approximately 10,000 neurons in its representative insect brain. Widely available experimental tools allow manipulation of gene expression and introduction of foreign transgenes in labeled neurons throughout the *Drosophila* brain, including in the learning and memory centers [9–12], whose synaptic connectivities can be reconstructed via an existing electron microscopy data set [10, 13, 14].

Larvae carry out complex behaviors including sensory-guided navigation [15–25], which can be modified by learning [3, 4, 6, 8]. Larval *Drosophila* has long been a model for the study of memory formation, with a well-established paradigm developed to study associative memory formation through classical conditioning [3, 4, 6–9, 26–28]. In this paradigm, larvae are trained and tested in groups, and learning is quantified by the difference in the olfactory preferences of differently trained groups of larvae. These assays quantify the effects of learning on a population level, but it is impossible to identify whether or to what extent an individual larva has learned.

New methods allow direct measurement of neural activity in behaving larvae [29–31] and reconstruction of the connections between the neurons in a larva’s brain [10, 13, 14, 32, 33], potentially allowing us to explore how learning changes the structure and function of this model nervous system. Using these tools requires us to identify larvae that have *definitively learned*. Recently, a device has been developed for assaying individual adult flies’ innate [34] and learned [35] olfactory preferences, but no comparable assay exists for the larval stage.

Further, to explore structural changes associated with learning, we need to form protein-synthesis dependent long-term memories [36–38]. Larvae trained to associate odor with electric shock form memories that persist for at least 8 hours [39]. Odor-salt memories have been shown to partially persist for at least 5 hours [40, 41] and can be protein-synthesis dependent [41], depending on the initial feeding state of the larva. Overnight memory retention, whether or not requiring protein-synthesis, has not been demonstrated in the larva, nor has long-lasting retention of appetitive memories.

In this work, we demonstrate a new apparatus for *in situ* training and measurement of olfactory preferences for individual larvae. We use this assay to quantify appetitive memories formed by presentation of Carbon Dioxide combined with optogenetic activation of reward neurons. Using this device, we find that larvae are sensitive to both the timing and context of the reward presentation, that learning is quantized and all-or-none, and that repeated presentation of odor without reinforcer can erase a newly formed memory. We induce memories that persist overnight, and control whether these memories require protein synthesis through alteration of the training protocol.

## Results

### A Y-maze assay to characterize olfactory preferences of individual *Drosophila* larvae

Establishing the degree to which an individual larva seeks out or avoids an odorant requires repeated measurements of that larva’s response to the odor. We developed a Y-maze assay [42] to repeatedly test an individual’s olfactory preference. The Y-mazes (**Figure 1**A) are constructed from agarose with channels slightly larger than than the larvae, allowing free crawling only in a straight line [43, 44]. An individual larva travels down one channel and approaches the intersection with the other two branches of the maze. Here the larva is presented with odorized air in one branch and pure air in the other. The larva then chooses and enters one of the two branches. When the larva reaches the end of its chosen channel, a circular chamber redirects it to return along the same channel to the intersection to make another choice. Custom computer vision software detects the motion of the larva while computer controlled valves manipulate the direction of airflow so that the larva is always presented with a fresh set of choices each time it approaches the intersection.

**Figure 1:**
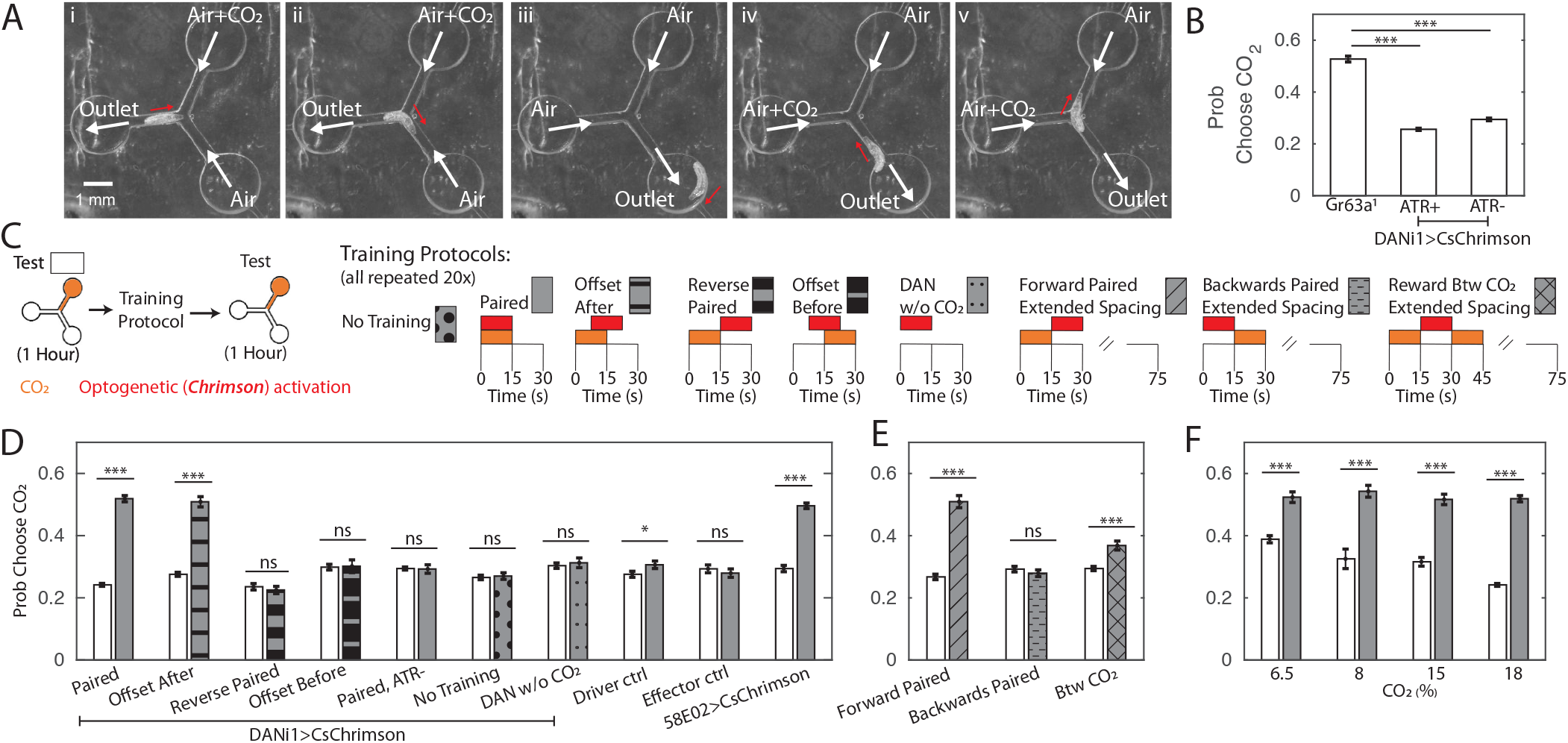
Y-maze assay to quantify innate and learned preference. (A) Image sequence of a larva making two consecutive decisions in the Y-maze assay. White arrows indicate direction of air flow; red arrow shows direction of larva’s head. (B) Probability of choosing channel containing CO_2_ without any training. (C) Schematic representation of experiments in (D,E,F). All larvae were tested in the Y-maze for one hour to determine initial preference and again following manipulation to determine a final preference. The manipulations were: Paired Training - reward in concert with CO_2_ presentation, 15s intervals, 20 repetitions; Offset After - reward presentation 7.5 s after CO_2_ onset, 15s intervals, 20 repetitions; Reverse-Paired Training - reward opposite CO_2_ presentation, 15s intervals, 20 repetitions; Offset Before - reward presentation 7.5 s before CO_2_ onset, 15s intervals, 20 repetitions; DAN Activation Without CO_2_ - CO_2_ is never presented, while reward is presented at 15s intervals, 20 repetitions; no training - no manipulation between two testing periods; Forward Paired (extended spacing) - 15s reward follows 15s CO_2_ presentation, followed by 60 seconds of air, 20 repetitions; Backwards Paired (extended spacing) - 15s reward prior to 15s CO_2_ presentation, followed by 60 seconds of air, 20 repetitions; Reward Between CO_2_ (extended spacing) - 15s reward presentation between two 15s CO_2_ presentations, followed by 45 seconds of air, 20 repetitions. (D) Probability of choosing CO_2_ containing channel before and after manipulation. All animals were fed ATR supplemented food, except those marked ATR-. (E) Probability of choosing CO_2_ containing channel before and after training as a function of reward timing, in training protocols with extended air spacings. All animals were DANi1>CsChrimson and fed ATR. (F) Probability of choosing CO_2_ containing channel before and after 20 cycles of paired training, as a function of CO_2_ concentration, used both during training and testing. All animals were DANi1>CsChrimson and fed ATR. * p<0.05, ** p<0.01, *** p<0.001.

We first sought to determine the suitability of this assay for measuring innate behavior. *Drosophila* larvae avoid Carbon Dioxide (CO_2_) at all concentrations [20, 45–48]. We presented larvae with a choice between humidified air and humidified air containing CO_2_ each time they approached the central junction. At the 18% concentration used throughout this work, larvae with functional CO_2_ receptors chose the CO_2_-containing channel about 25% of the time. The probability of choosing the CO_2_ containing channel decreased as CO_2_ concentration in that channel increased (**Figure 1**F). *Gr63a*^1^ [46] larvae lacking a functional CO_2_ receptor were indifferent to the presence of CO_2_ in the channel (**Figure 1**B), indicating that larvae responded to the presence of CO_2_ and not some other property of the CO_2_ containing air stream.

### Pairing CO_2_ presentation with optogenetic activation of a single pair of reward neurons eliminates CO_2_ avoidance

Activation of the DAN-i1 pair of mushroom body input neurons has been shown to act as a reward for associative learning [9, 28, 49, 50]. In previous experiments, the conditioned odor was innately attractive. We wondered whether pairing DAN-i1 activation with CO_2_ would lessen or even reverse the larva’s innate avoidance of CO_2_.

To train larvae in the same Y-maze used to measure preference, we manipulated the valves so that the entire chamber was either filled with humidified air or with humidified air mixed with additional CO_2_, independent of the position of the larva, which was not tracked during training. At the same time, we activated DAN-i1 neurons expressing CsChrimson using red LEDs built in to the apparatus. For some larvae, we activated DAN-i1 when CO_2_ was present (paired, **Figure 1**D). For others, we activated the reward neurons when only air was present (reverse-paired, **Figure 1**D). Each training cycle consisted of one 15 second air presentation and one 15 second CO_2_ presentation, with DAN-i1 activated for the entirety of the air (reverse-paired) or CO_2_ (paired) presentation phase. For each larva, we first measured naive preference and then preference following training.

Note that the training protocols schematized in **Figure 1**D were repeated for 20 successive cycles. Thus, for instance, in the reverse-paired scheme CO_2_ offset at t=15s coincided with reward onset, and the reward offset at t=30s coincided with CO_2_ onset at t=0 of the subsequent cycle.

We found that in the paired group, larvae became indifferent to CO_2_ presentation following 20 training cycles (**Figure 1**D, DANi1*>*CsChrimson, Paired). We did not observe any change in preference in the reverse-paired group (DANi1*>*CsChrimson, Reverse-Paired). Nor did we observe a preference change following paired training for genetically identical animals not fed all-*trans*-retinal (ATR), a necessary co-factor for CsChrimson function (DANi1*>*CsChrimson, Paired ATR-). Animals fed ATR but not exposed to red light failed to show a preference shift (DANi1*>*CsChrimson, No Training). Larvae of the parent strains fed ATR and given paired training showed no preference shift (Effector Control, Driver Control). To control for possible effects of DAN-i1 activation, we activated DAN-i1 in 15 second intervals without presenting CO_2_ at all during the training (DANi1*>*CsChrimson, DAN w/o CO_2_); these larvae showed no shift in preference.

Taken together these results show that the change in CO_2_ preference requires activation of the DAN-i1 neurons and is not due to habituation [51–53], red light presentation, or other aspects of the training protocol. In particular, the paired and reverse-paired group experienced identical CO_2_ presentations and DAN-i1 activations with the only difference the relative timing between CO_2_ presentation and DAN-i1 activation.

Activation of DAN-i1 coincident with CO_2_ presentation decreased larvae’s subsequent avoidance of CO_2_. Formally, this admits two possibilities: the larva’s preference for CO_2_ increased because CO_2_ was presented at the same time as the reward or because CO_2_ predicted the reward. To test whether learning was contingent on coincidence or prediction, we carried out an additional set of experiments. As before, we first tested innate preference, then presented 20 alternating cycles of 15s of CO_2_ followed by 15s of air. However, this time during the conditioning phase, we either activated DAN-i1 7.5 seconds *after* CO_2_ onset, in which case CO_2_ predicted DAN-i1 activation, or 7.5 seconds *before* CO_2_ onset, in which case CO_2_ predicted withdrawal of DAN-i1 activation.

In both cases DAN-i1 was activated in the presence of CO_2_ for 7.5 s and in the presence of air alone for 7.5 s. If learning depended only on the coincidence between reward and CO_2_ presentations, both should be equally effective at generating a change in preference. In fact, we only found an increase in CO_2_ preference following training in which the CO_2_ predicted the reward (**Figure 1**D).

Next we asked whether reward prediction alone was sufficient to establish a memory, or if coincidence between CO_2_ and DAN-i1 activation was also required. We altered the training protocol to present 15 seconds of CO_2_ followed by 60 seconds of air. Some larvae were rewarded by activation of DAN-i1 in the 15 seconds immediately following CO_2_ presentation (Forward Paired), while others were rewarded in the 15 seconds immediately prior to CO_2_ presentation (Backwards Paired). For a third group of larvae, CO_2_ was presented both before and after reward presentation (reward between CO_2_ presentations). At no time was DAN-i1 activated in the presence of CO_2_, but in the first group CO_2_ predicted DAN-i1 activation while in the others it did not. We found an increased CO_2_ preference for animals in this first group only (**Figure 1**E), indicating that reward prediction is both necessary and sufficient for learning in this assay.

In other associative conditioning experiments using DAN-i1 activation as a reward, decreased attraction to the odor was observed in the reverse-paired groups [9, 49, 50]. In our experiments, we did not see any evidence of increased aversion in the reverse-paired groups.

We wondered whether it might be possible to achieve an attraction to CO_2_ following training, rather than ‘merely’ a loss of avoidance. In other contexts, reward via activation of 3 DANs (DAN-i1, DAN-h1, DAN-j1 - whether DAN-h1 is present in second instar larvae, used in this study, is presently unreported) labeled by the 58E02-Gal4 line has been reported to produce strong learning scores [9, 50, 54, 55]. We repeated the training protocol, substituting 58E02 activation for DAN-i1 activation alone, but did not see an increased preference following training compared to DAN-i1 activation alone (**Figure 1**D, 58E02*>*CsChrimson).

Next we asked how varying the CO_2_ concentration might affect animals’ performance in the assay. We presented lower concentrations of CO_2_ both during the training and testing phases, and found that decreasing the CO_2_ concentration decreased innate avoidance of CO_2_. In all cases, following training, larvae lost avoidance to CO_2_ but none showed statistically significant attraction (**Figure 1**F).

### Learning is quantized and all-or-none

We investigated how change in preference for CO_2_ following associative conditioning with DAN-i1 activation depended on the amount of training. As in the previous experiments, we first measured the innate preference, then trained the larva using repeated cycles alternating pure and CO_2_ containing air, while activating DAN-i1 in concert with CO_2_ presentation. In these experiments however, we varied the number of training cycles an individual larva experienced. We found that as a group, larvae that had experienced more training chose the CO_2_ containing channel more often (**Figure 2**A).

**Figure 2:**
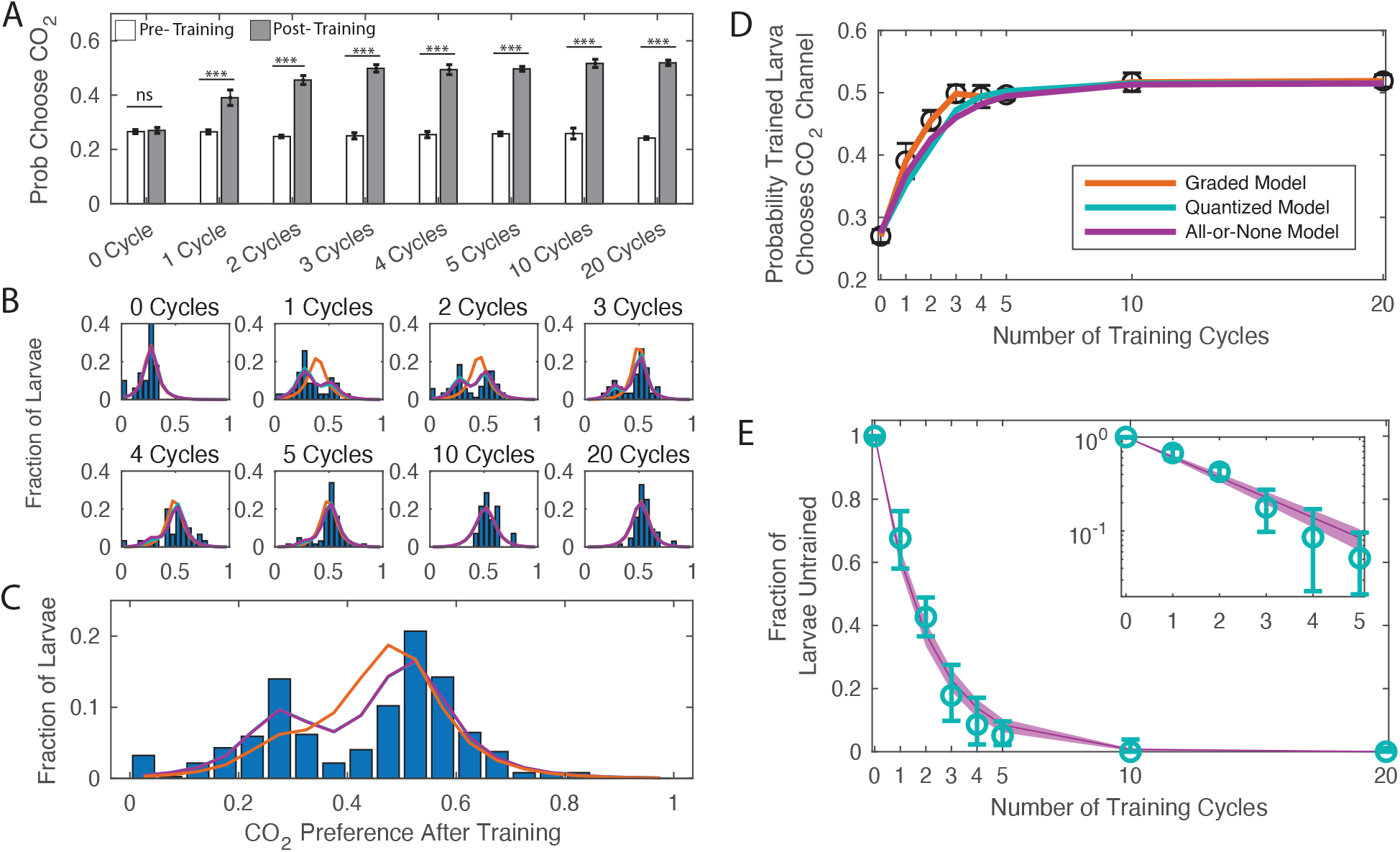
Dose dependence of learning. DANi1>CsChrimson were given varying cycles of paired training (as in **Figure 1**C). (A) Probability of choosing CO_2_ containing channel before and after training, as a function of amount of training. *** p<0.001. (B) Histograms of individual larva preferences after training, grouped by number of training cycles. (C) Histogram of individual larva preference after training for all larvae. (D) Population average probability of choosing CO_2_ following training vs. dose. (E) Fraction of larvae untrained vs. number of training cycles. Teal: fit parameters and error ranges from quantized model, purple lines, prediction and error ranges from memoryless model. Note logarithmic y-axis on insert. (C-E) Orange: graded model prediction - post-training preference is represented by a single Gaussian distribution whose mean and variance depend on amount of training; Teal: quantized model prediction - post-training preference is represented by two fixed Gaussian distributions and the fraction of larvae in each population depends on the amount of training; Purple: all-or-none model prediction - post-training preference is represented by two fixed Gaussian distributions and the effect of a single training cycle is to train a fixed fraction of the remaining untrained larvae.

Our data showed that increasing the amount of training increased overall preference for CO_2_ up to a saturation point. But what was the mechanism for this change? Did each cycle of training increase each larva’s preference for CO_2_ by some small amount, with the effect accumulating over repeated training (graded learning)? Or did some larvae experience a dramatic preference change - from naive to fully trained - with each cycle of training, with the number of fully trained larvae increasing with training repetitions (quantized learning)?

Either quantized or graded learning can explain the shift of mean preference of a population; to differentiate between the modes of learning, we examined repeated decisions made by individual animals, measurements that were impossible in previous larval assays. For each larva, we quantified the change in CO_2_ preference before and after training. **Figure 2**B shows a histogram of larva preference (the fraction of times an individual larva chose the CO_2_ containing channel) after training, grouped by the number of cycles of training a larva received.

Larvae that received no training (0 cycles) formed a single population that chose CO_2_ 27% of the time. Larvae that were trained to saturation (20 training cycles) also formed a single group centered around 52% probability of choosing CO_2_. Both the graded and quantized learning models make the same predictions for these endpoints, but their predictions vary starkly for the intermediate cases. A graded learning model predicts that all larvae that received the same amount of training would form a single group whose mean preference for CO_2_ would increase with increasing training. A quantized learning model predicts that larvae that have received the same amount of training will form two discrete groups (‘trained’ and ‘untrained’) with fixed centers whose means do not depend on the amount of training. With increased training an increasing fraction of larvae would be found in the trained group.

We fit the distributions of preference following conditioning to graded and quantized learning models. In the graded model, the preference was represented by a single Gaussian distribution whose mean and variance were a function of amount of training (orange, **Figure 2**). In the quantized model, the preference was represented by two Gaussian distributions; the fraction of larvae in each population was a function of the amount of training (teal, **Figure 2**).

We found that the data were better described by the quantized learning model (**Table S3**): larvae form two discrete groups, with the fraction in the trained group increasing with each cycle of additional training. The centers of the two groups do not vary with the amount of training, a point made most clear by considering the preference after training of all larvae taken together regardless of the amount of training received (**Figure 2**C), which shows two well defined and separated groups. From these data, we concluded that the effect of our associative conditioning on an individual larva is to either cause a discrete switch in preference or to leave the initial preference intact.

Next we asked what effect, if any, associative conditioning had on larvae that retained their innate preferences following training. Whether humans form associative memories gradually through repeated training or learn in an all-or-none manner has been the subject of debate in the Psychology literature [56]; recent electrophysiological measurements in humans supports the all-or-none hypothesis [57]. Our fit to the quantized learning model produces an estimate of the fraction of larvae that remain untrained following training. If the training were *cumulative*, we would expect a threshold effect: as the number of cycles of training increased from 0, most larvae would initially remain untrained until a critical number of cycles were reached and there would be a sudden shift to a mostly trained population. But when we plotted the fraction of untrained larvae vs. number of training cycles, we saw that the fraction of larvae in the untrained group exponentially decreased with increasing training (**Figure 2**E, note logarithmic y-axis on insert). Exponential decay is an indicator of an all-or-none process - rather than accumulating over time, each training cycle either caused a larva to switch preference entirely or had no effect.

We confirmed this interpretation by fitting the population distributions to an all-or-none quantized learning model in which the effect of a single training cycle was to train a fixed fraction of the remaining untrained larvae (purple, **Figure 2**). This model fit the data better than the graded learning model and almost as well as the original quantized learning model (in which the fraction of untrained larvae was fit separately to each group) despite having fewer parameters than either model. According to standard selection rules (BIC and AIC), the all-or-none quantized model best describes the data (**Table S3**).

### Repeated exposure without reward following training leads to memory extinction; overnight consolidation makes memories resistant to extinction

Reversal learning, in which the reward contingency is reversed, and extinction, in which the conditioned stimulus is presented without reward, experiments explore cognitive flexibility. Previous experiments with both adult *Drosophila* [58–61] and larval [62] *Drosophila* demonstrated a reversal learning paradigm. Extinction has been demonstrated in adult flies [63–65] but not in larvae.

To test for extinction, we again first measured an individual larva’s CO_2_ preference and then carried out associative conditioning for a given number (2- of training cycles. Next instead of immediately testing the larva’s new preference for CO_2_, we exposed the larva to an extinction phase - 18 cycles of alternating CO_2_ and air without any optogenetic reward. Following the extinction period, we tested larvae as usual to measure their changed preference for CO_2_. As a control against the effects of increased CO_2_ exposure, we also performed habituation experiments, which were the same as the extinction experiments, except the 18 unrewarded cycles were presented *prior* to the rewarded training cycles. The extinction and habituation protocols are schematized in **Figure 3**A.

**Figure 3:**
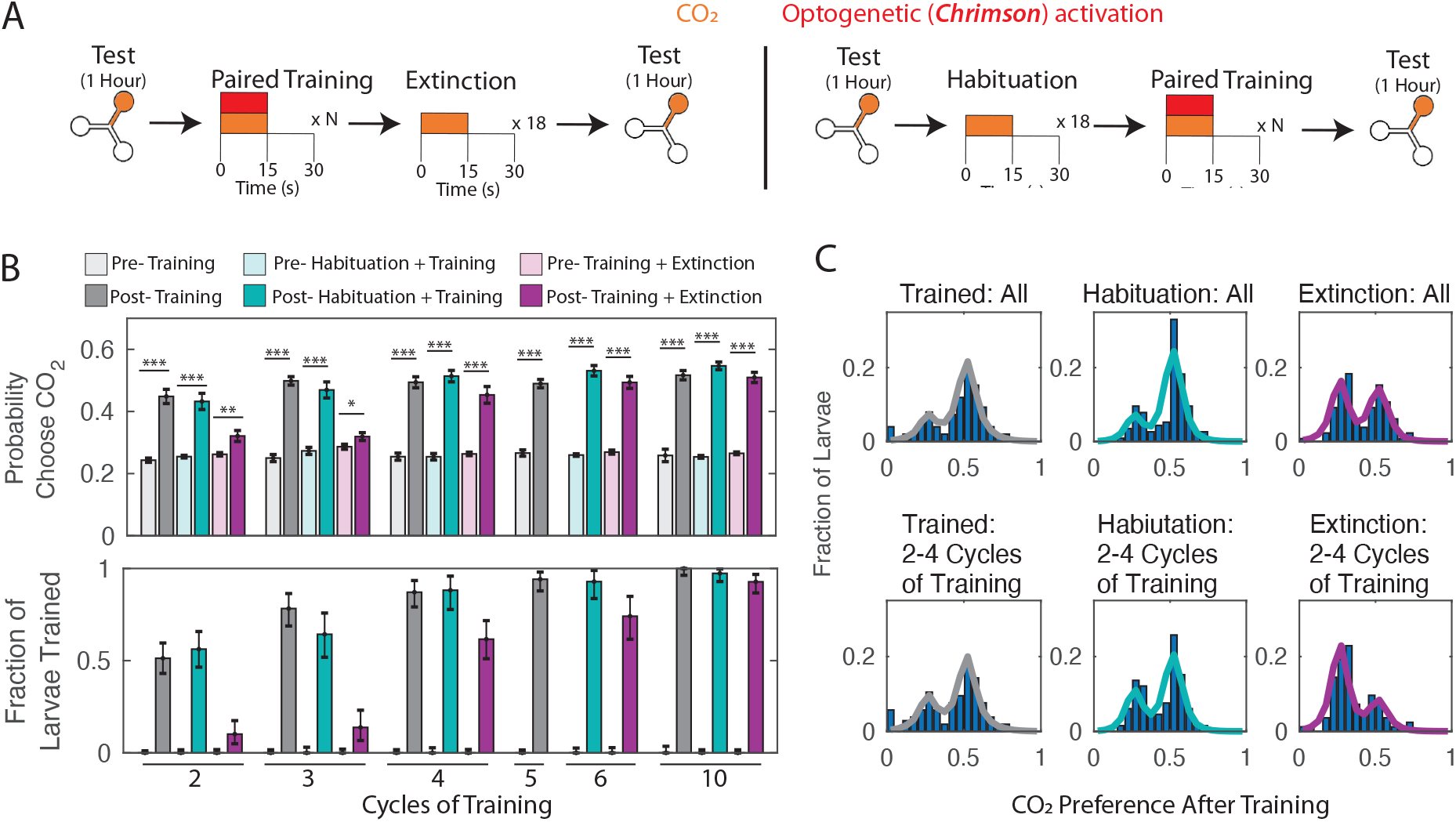
Memory extinction. (A) Testing and training protocols for B,C. Training + Extinction: larvae were exposed to 18 cycles of alternating CO_2_ and air following training. Habituation + Training: larvae were exposed to 18 cycles of alternating CO_2_ and air prior to training. (B) Probability of choosing CO_2_ containing channel (top) and fraction of larvae in trained group according to double Gaussian model fit (bottom) before and after training scheme. (C) Histograms of individual larva preference after training, for all larva and for larva trained with 2-4 training cycles. * p<0.05, ** p<0.01, *** p<0.001.

When we compared the ‘habituated’ groups of larvae to larvae trained for the same number of cycles without habituation or extinction, we found that un-rewarded CO_2_ presentation prior to training had no effect on the eventual preference change (**Figure 3**B). This was unsurprising, as the initial testing period already offered a number of unrewarded CO_2_ presentations. In contrast, unrewarded CO_2_ presentations *following* training reversed the effect of training; for small (2 or 3 cycles) amounts of training, the reversal was almost complete (**Figure 3**B).

We previously observed that associative conditioning produced a discrete and quantized change in CO_2_ preference. Here we found that extinction following training greatly reversed the effects of conditioning. We wondered whether larvae that had been subject to both training and extinction reverted to their original CO_2_ preference or to an intermediate state. In the former case, we would expect to see a bimodal distribution of preference change following training and extinction, while in the latter we would see a third group of larvae. This group would be most evident in experiments where 2-4 cycles of training were followed by extinction, as these had the largest deficit in the fraction of trained larvae compared to habituated larvae that received the same amount of training. We examined the preferences of all larvae following 2-4 cycles of training, grouped by whether they were normally trained, habituated, or subject to extinction (**Figure 3**C). In all cases, we observed two groups with the same central means and no evidence of a third intermediate group. We concluded that larvae subject to training then extinction reverted to their “untrained” state.

### Larvae can retain memory overnight

Studies in adult [36, 58, 66] and larval [39–41, 67–69] *Drosophila* have identified distinct memory phases: short-term memory (STM), middle-term memory (MTM), long-term memory (LTM) and anesthesia-resistant memory (ARM). LTM and ARM are both consolidated forms of memory, which are thought to be represented by different, separate pathways [70]. ARM is resistant to anesthetic agents [71]; LTM requires cAMP response element-binding protein (CREB) dependent transcription and *de-novo* protein synthesis, while ARM does not [36, 38]. Adults have been shown to retain memories for up to a week [36]. Larvae trained to associate odor with electric shock form memories that persist for at least 8 hours [39]. Odor-salt memories have been shown to persist for at least 5 hours [40, 41] and can be either ARM or LTM, depending on the initial feeding state of the larva.

We sought to determine whether we could create consolidated memories that would persist overnight, and if so, whether these memories represented ARM or LTM.

As in the previously described experiments, we first tested each larva’s individual preference in the Y-maze assay, trained it to associate CO_2_ presentation with DAN-i1 activation, and then measured its individual preference again following training. After this second round of testing, we removed the larva from the apparatus and placed it on food (without ATR) overnight. The next day we placed the larva back in the Y-maze and again tested its preference for CO_2_, without any additional training.

We found that following twenty cycles of training, larvae became indifferent to CO_2_ and this in-difference persisted to the next day. Similarly, we found that most larvae switched preference following five cycles of training and retained that preference overnight. Larvae that received no training or 20 cycles of unpaired training had no change in CO_2_ preference immediately following training or the next day (**Figure 4**B).

**Figure 4:**
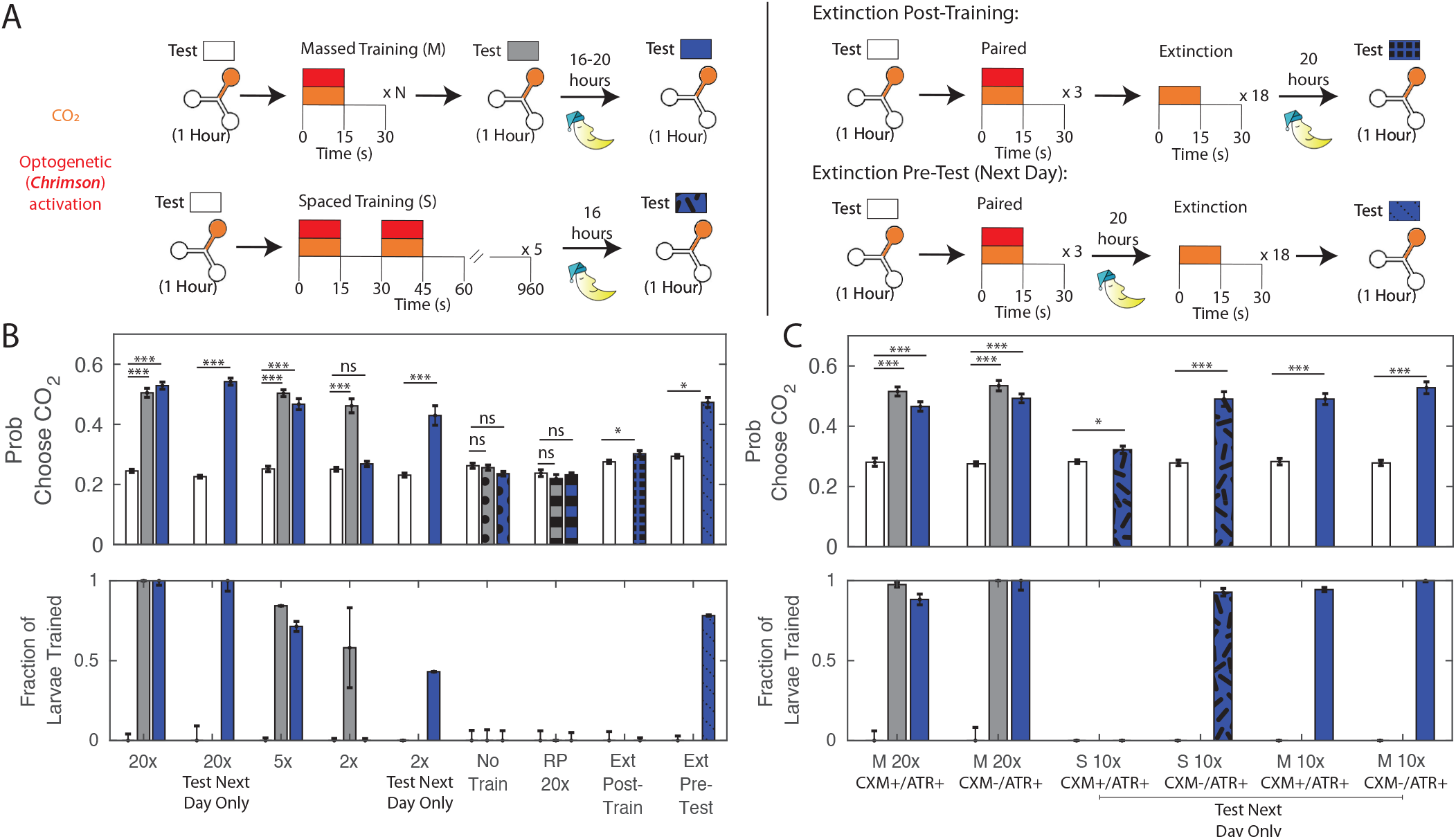
Memory retention overnight. (A) Testing and training protocols. Larvae were tested, trained immediately after testing, tested again, then placed on food overnight and tested the following day. For extinction experiments, larvae were trained 3 times, and then exposed to 18 cycles of alternating CO_2_ and air either immediately following training or prior to testing the next day. All training was massed unless otherwise indicated. All larvae were DANi1>CsChrimson. Larvae were raised on food containing ATR, except for ATR+/CXM-, ATR+/CXM+ larvae who were fed ATR supplemented yeast paste (without/with cycloheximide) for 4 hours prior to initial testing. For reverse-paired (RP) and no training schemes, see **Figure 1**B. (B,C) Probability of choosing CO_2_ containing channel (top) and fraction of larvae in trained group according to double Gaussian model fit (bottom) prior to training, immediately following training, and the next day. When the center bar is missing, larvae were not tested immediately following training but instead removed immediately to food. * p<0.05, ** p<0.01, *** p<0.001.

We had previously shown two cycles of training caused roughly half the larva to change preference immediately after training. We decided to use this partition to verify a correlation between immediate and long-term memories; we expected that larvae initially in the ‘trained’ group would also form a ‘trained’ group the following day. However, while we found that two cycles of training were sufficient to cause some larvae to become indifferent to CO_2_ immediately following training, when we tested these larvae the next day, we found that all had reverted to their initial avoidance of CO_2_.

There were two possible explanations for this reversion. Perhaps two cycles of training were sufficient to form a short term memory, but more training was required to induce a long-term memory. Or perhaps the *testing* period, in which larvae were exposed repeatedly to CO_2_ without reward, reversed the two-cycle training. To control for the latter, we modified the experimental protocol. We tested each larva’s innate preference, presented two training cycles, and then immediately removed the larva to food overnight, without any further testing. When we tested these larvae the next day, we found that they showed decreased avoidance of CO_2_. This indicated that two cycles of training were sufficient to form a memory lasting overnight, but that immediate exposure to unrewarded CO_2_ following this short training interval likely reversed the effects of training, an effect we observed in **Figure 3**. When larvae were trained for 20 cycles, omitting the testing had no effect on these larvae’s preferences the following day.

To confirm that extinction could explain the failure to form a persistent memory, we exposed larvae to three cycles of paired training, then 18 cycles of extinction (as in **Figure 3**) and then removed them to food overnight before testing their preferences the next day. As expected, these larvae avoided CO_2_ as much the next day as they did prior to training (**Figure 4**B, Ext Post-Train).

We wondered whether memories that had consolidated overnight would be more resistant to extinction. We repeated the previous experiment with a single modification. As before, we tested the larva’s initial preference and trained it with three cycles of rewarded CO_2_ presentation. This time, we immediately removed the larva to food following training. The next day, we returned the larva to the Y-maze and presented the extinction phase of 18 unrewarded CO_2_ presentations prior to testing for CO_2_ preference. We found that in this case, larvae still expressed an increased preference for CO_2_ despite the extinction phase (**Figure 4**B, Ext Pre-Test). The only difference between the two experiments was whether we attempted extinction immediately after training or the next day. Thus we concluded that overnight consolidation made memories more resistant to extinction.

ARM can be distinguished from LTM because the latter requires *de novo* protein synthesis and can be disrupted by ingestion of the translation-inhibitor cycloheximide (CXM). To incorporate CXM feeding, we modified our protocols. Instead of raising larvae on ATR supplemented food, we raised them on standard food and then fed them with ATR supplemented yeast paste for 4 hours prior to the experiment (ATR+/CXM-). For some larvae (ATR+/CXM+), we also added CXM to the yeast paste. In this way, we could be sure that if ATR+/CXM+ larvae ingested enough ATR to allow for CsChrimson activation of DAN-i1, they must have also ingested CXM as well. To further verify CXM ingestion, we placed ATR+/CXM+ and ATR+/CXM-larvae on clean food and allowed them to continue development. 95% of ATR+/CXM-larvae pupated, while only 45% of ATR+/CXM+ larvae pupated.

Following the 4 hour feeding period, ATR+/CXM+ and ATR+/CXM-larvae were treated identically. As in the previously described experiments, we first tested each larva’s individual preference in the Y-maze assay, trained the larva twenty times to associate CO_2_ presentation with DAN-i1 activation, and then measured its individual preference again following training. After this second round of testing, we removed the larva from the apparatus and placed it on food (without ATR or CXM) overnight. The next day we placed the larva back in the Y-maze and again tested its preference for CO_2_, without any additional training.

We found that performances tested immediately and 16 hours after training were both unaffected by CXM treatment. Following twenty cycles of training, larvae from both groups (ATR+/CXM+; ATR+/CXM-) became indifferent to CO_2_ and this indifference persisted to the next day (**Figure 4**C). This suggests that the memory formation was independent of *de novo* protein synthesis.

In adult *Drosophila*, whether ARM or LTM is formed depends on the training protocol [36, 58, 72–74]. ‘Massed’ training, in which all conditioning occurs in rapid sequence without rest intervals, results in ARM, while ‘spaced’ training, in which the conditioning occurs in blocks separated by intervals of time, produces LTM. Our training protocol more closely resembles massed training, so it seemed sensible that it would produce ARM. To see if we could also develop LTM, we established a spaced training protocol. Larvae received two paired cycles of training, followed by a 15 minutes interval of air-presentation only; this sequence was repeated five times. To keep the total length of the experiment within a (covid-related) limited daily time window, we did not test the larvae immediately after training but only the next day.

Prior to spaced training, both ATR+/CXM- and ATR+/CXM+ larvae avoided CO_2_ to the same degree. We found that one day following spaced training, ATR+/CXM+ larvae continued to avoid CO_2_, while ATR+/CXM-larvae did not. This indicated that spaced training formed a memory whose retention was disrupted by CXM. To verify that spacing the trials was essential to forming a protein-synthesis dependent memory, we duplicated the experiments exactly, except we presented 10 cycles of training *en masse*, rather than spacing them. In this case, both ATR+/CXM- and ATR+/CXM-larvae expressed learned indifference to CO_2_ one day following training (**Figure 4**C).

## Discussion

In this work, we demonstrated a new apparatus for training individual larvae and assessing their olfactory preferences. Compared to the existing paradigm, our assay allows for measuring individual animals’ changes in preference due to training, allows for greater control of the temporal relation between the conditioned and unconditioned stimuli, and does not require any handling of the animals between training and testing.

In our assay, larvae learned in a switch-like (all-or-none two-state quantized) manner. The learning process was better described as a sudden transition between states rather than a graded change in preference. We found that each cycle of training (presentation of CO_2_ coupled with reward) either caused a state transition or did not; we found no evidence of a cumulative effect of prior training in the probability that a given cycle of training would induce a state transition in larvae that had not already transitioned. We did however find evidence that repeated cycles of training stabilized memories against later extinction effected by presentation of CO_2_ without reward. These measurements were enabled by our assay’s ability to track individual preferences over the course of the entire experiment.

We found that larvae trained in our assay retained memories overnight: 16-20 hours. When training was presented all at once, these memories were not disrupted by ingestion of the protein-synthesis inhibitor cycloheximide, while when training was spaced over time, cycloheximide feeding prevented long duration memory formation. Thus we identified spaced training as producing long term memory (LTM) and the massed training as producing anaesthesia resistant memory (ARM). These results are the first demonstrations that larvae can retain memories overnight; they are entirely congruent with observations in adult flies.

We explored how the order of CO_2_ and reward presentations affected learning. We found that for larvae to learn CO_2_ onset must occur coincident with or before reward onset, but that it was neither necessary nor sufficient for CO_2_ and reward to be presented together at the same time. Here our results differed from previous reports. In other assays, presenting the reward (including via activation of DAN-i1) prior to presenting the conditioned odor results in *decreased* attraction/increased avoidance [9, 50] of that odor. We found that such “reverse-pairings” neither increased nor decreased a larva’s avoidance of CO_2_. There are a number of differences - our new behavioral assay, our use of the innately aversive CO_2_ as the conditioned odor, our activation of DAN-i1 via CsChrimson rather than ChR2-XXL, our focus on second rather than third instar larvae - that might account for the discrepancy.

One intriguing hypothesis is that this difference might be encoded in the connectivities of neurons representing CO_2_ in the mushroom body. In the adult fly innate CO_2_ avoidance requires MBON output [75, 76]. As in the adult [77–81], larval MBONs encode either approach or avoidance and synapse onto a convergence neuron that integrates their activities [82]. Prior to training, the approach and avoidance pathways are thought to be in balance. Learning that a stimulus is appetitive *weakens* the connection between KCs encoding that stimulus and the *avoidance* MBONs, promoting approach, while aversive conditioning weakens the connection between KCs and approach MBONs. Our observations that appetitive conditioning can eliminate CO_2_ avoidance but not promote CO_2_ approach and that CO_2_ avoidance cannot be increased by reversing the reward contingency would both be explained if there are no connections from neurons representing CO_2_ to approach promoting MBONs. In this case, the only effect of learning would be to decrease/eliminate the ability of CO_2_ to provoke an avoidance response.

Understanding memory formation at the circuit and synaptic levels simultaneously is a heroic task, even aided by the larva’s numerically simple nervous system and the tools (including EM-reconstruction) available in the larva. The work here represents progress towards this goal. We demonstrate long-term protein synthesis dependent memory, implying that memories are encoded in synaptic change. Our assay allows us to precisely identify those individuals who have formed long-term memories. Animals are found in only two behavioral states: innate avoidance or learned indifference; this likely reflects two discrete states of the underlying neural circuit.

That there are only two behavioral states and that associative conditioning produces indifference rather than attraction to CO_2_ is most parsimoniously explained by the circuit mediating innate CO_2_ avoidance passing through a bottleneck that is gated down-stream of DAN-i1. Even if the circuit is not this simple, it remains particularly favorable to analysis. The conditioned stimulus is sensed by a single pair of genetically identified sensory neurons; the unconditioned stimulus is provided by activation of a single pair of genetically identified reward neurons whose connectivity has been fully reconstructed [50]. How the larva navigates in response to CO_2_ presentation has been described in detail [20, 45, 48, 83], as has been how neurons downstream of DAN-i1 and the KCs contribute to navigational decision making [9, 10, 49, 50]. This is a particularly favorable starting point to understand how synaptic plasticity due to associative conditioning leads to changes in circuit function that effect changed behavioral outcomes.

## Conclusion

We introduced a Y-maze assay capable of measuring the olfactory preferences of individual larval *Drosophila* and of *in situ* associative conditioning. We found that when larvae learn to associate CO_2_ with reward neuron activation, the result is a switch from innate avoidance to learned indifference, with no intervening states. We demonstrated a protocol to form stable protein-synthesis dependent long term memories. This provides a strong starting point for ‘cracking’ a complete olfactory learning circuit.

## Acknowledgements

This project was supported by NSF grant 1455015, NIH grant DP2-EB022359, and a Sloan Foundation fellowship to MHG. The funders had no role in the design or analysis of the experiments.

The following ORCIDs apply to the authors: 0000-0001-6611-5941 (AL), and 0000-0001-7528-6101 (MG).

## Author Contributions

AL; Conceptualization, Methodology, Data curation, Software, Formal analysis, Investigation, Writing — original draft, Visualization, Writing — review and editing; JT: Software; JW: Investigation; MG: Conceptualization, Methodology, Formal analysis, Supervision, Funding acquisition, Writing — original draft, Project administration, Visualization, Writing — review and editing

## Materials and Methods

### Fly strains

The following fly strains were used:

- 20XUAS-CsChrimson-mVenus (Bloomington Stock #55136)
- SS00864 split-Gal4 (gift from Marta Zlatic, Janelia Research Campus)
- w[*]; Gr63a[1] (Bloomington Stock #9941)
- w[1118]; Py[+t7.7] w[+mC]=GMR58E02-GAL4attP2 (Bloomington Stock #41347)

### Crosses

Virgin female *UAS-CsChrimson* flies were crossed with males of the split-Gal4 driver strain SS00864-Gal4.

For experiments in **Figure 1**D (R58E02*>*CsChrimson, Paired Training), virgin female *UAS-CsChrimson* flies were crossed with males of the Gal4 driver strain GMR58E02-GAL4.

### Larva collection

Flies were placed in 60 mm embryo-collection cages (59-100, Genessee Scientific) and allowed to lay eggs for 6 hours at 25C on enriched food media (Nutri-Fly German Food, Genesee Scientific). For all experiments except otherwise specified, the food was supplemented with 0.1 mM all-trans-retinal (ATR, Sigma Aldrich R2500). Cages were kept in the dark during egg laying. When eggs were not being collected for experiments, flies were kept on plain food at 18C.

Petri dishes containing eggs and larvae were kept at 25C in the dark for 48-60 hours. Second instar larvae were separated from food using 30% sucrose solution and washed in water. Larvae were selected for size. Preparations for experiments were carried out in a dark room.

### Y-maze

We used SLA three-dimensional printing to create microfluidic masters for casting [29, 84]. Masters were designed in Autodesk Inventor and printed on an Ember three-dimensonal printer (Autodesk) using black prototyping resin (Colorado Photopolymer Solutions). After printing, masters were washed in isopropyl alcohol, air-dried, and baked at 65C for 45 minutes to remove volatile additives and non-crosslinked resin. 4% agarose (Apex Quick Dissolve LE Agarose, Cat #20-102QD, Genesee Scientific) was poured over the masters and allowed to solidify; then mazes were removed from the mold. Agarose Y-mazes were stored in tap water before use.

The mazes are 1 mm in depth. Each channel is 1.818 mm in length and 0.4 mm in width, and ends in a circular chamber (radius = 1 mm) which redirects larva back to the intersection. An inlet channel (depth = 0.1 mm, length = 1.524 mm, width = 0.1 mm) to the circular chamber connects to tubing for our network of air, CO_2_, and vacuum sources.

### Behavioral experiments

Individual larvae were selected for size and placed into a Y-maze using a paintbrush. The Y-maze was placed into a PDMS (Sylgard 184, 10:1 base:curing agent) base, where tubing was secured. The Y-maze and base were encased in a dark custom-built box. Larvae were monitored under 850 nm infrared illumination (Everlight Electronics Co Ltd, HIR11-21C/L11/TR8) using a Raspberry Pi NoIR camera (Adafruit, 3100), connected to a Raspberry Pi microcomputer (Raspberry Pi 3 Model B+, Adafruit, 3775). Experiments were recorded using the same camera, operating at 20 fps. Eight copies of the assay were built, to assay the behaviors of multiple larvae in parallel.

Pressure for air, CO_2_, and vacuum were controlled at the sources (for vacuum regulation: 41585K43, McMaster-Carr; for pressure regulation: 43275K16, McMaster-Carr). CO_2_ and air were humidified through a bubble humidifier. Vacuum, air, and CO_2_ tubing to individual assays were separated through a block manifold after pressure control and humidification (BHH2-12, Clippard).

The CO_2_ concentration in the odorized channels was controlled by a resistive network of tubing connected to the air and odor sources. This inexpensive alternative to a mass-flow controller produced a stable ratio of odor to air that was consistent from day to day and independent of the overall flow rate. The direction of flow was controlled by solenoid pinch valves (NPV2-1C-03-12, Clippard), actuated by a custom circuit we designed.

Custom computer vision software detected the location of the larva in real time. Based on the larva’s location, computer controlled valves manipulated the direction of airflow so that the larva was always presented with a fresh set of choices each time it approached the intersection. The software randomly decided which channel would contain air and which contained air mixed with CO_2_.

In each maze, one channel was selected to be the outlet for flow and the other two were inlets. An individual larva began in the outlet channel and approached the intersection of the Y-maze, then chose to enter either an inlet branch containing air with CO_2_ or an inlet branch containing air only. When the larva’s full body entered the chosen channel, software recorded the larva’s choice of channel. When the larva reached the end of that channel and entered the circular chamber, valves switched to turn off CO_2_ and to switch vacuum to the channel containing larva, making that channel the outlet. The CO_2_ remained off (the larva experienced only pure air) until the larva exited the circular chamber. When the larva exited the circular chamber and proceeded towards the intersection, CO_2_ was introduced to one randomly selected inlet channel.

Software recorded the location of the larva at every frame (approx 20 Hz); the direction of airflow in the maze (which channel(s) had air; which channel had CO_2_ mixed with air, if any; and which channel had vacuum); and all decisions the larva made. We recorded when larvae entered or left a channel, and whether that channel presented CO_2_. Larvae could take three actions as they approached the intersection: choose the channel containing air with CO_2_ (scored as APPROACH); choose the channel containing pure air (scored as AVOID); or move backwards into their original channel before they reach the intersection. If a larva backed up and reentered the circular chamber it departed from before reaching the intersection, the software reset and presented the larva with a fresh set of choices when it next left the circle. We did not score backing up as a choice of either CO_2_ or air.

Following an hour of testing, larvae were trained in the same Y-maze assay used to measure preference. During the training period, unless described otherwise, each 30-second training cycles alternated 15 seconds of CO_2_ presentation, where both inlet channels contained a humidified mix of CO_2_ and air; followed by 15 seconds of air presentation, where both inlet channels had humidified air alone. This cycle was repeated some number of times (specified for each experiment in the figures). Red LEDs (Sun LD, XZM2ACR55W-3) integrated into the setup were used to activate CsChrimson synchronously with CO_2_ presentation (paired) or air presentation (reverse-paired).

The volume of the flow chamber was 11.68 mm^3^ and the volume of the tubing downstream of the valves is approximately 214 mm^3^. The flow rate exceeded 560 mm^3^/second, and the state of the chamber was taken to be the same as the state of valves.

Following training, larvae were tested for one hour in an identical scheme to that previously described for the naive measurement.

After larvae were placed into the Y-maze, larva were left in the maze in the dark for a minimum of 5 minutes. If a larva was not moving through the maze after 5 minutes, the larva was replaced before the experiment began. If larvae stopped moving through the maze during the first hour of testing, larvae were removed from the maze before training and results were discarded. This happened infrequently (approximately 5% of experiments).

### Overnight memory formation

For the memory retention experiments of **Figure 4**, testing and training followed identical procedures as above to establish larva preference. After the second round of testing testing, the larvae were removed from the Y-maze assay with a paintbrush and transferred to an individual 4% agar plate (30 mm, FB0875711YZ Fisher Scientific), with yeast paste added. Larva were kept in the dark at 18 C for approximately 20 hours. Prior to experiments the next day, larva were removed from the agar plate and washed in water before being placed in a new Y-maze. Larvae were then tested for CO_2_ preference for one hour as previously described. In all experiments in which larvae were removed from the apparatus and later retested, they were placed in the same apparatus but with a new agar Y-maze. Out of 317 larvae placed on agar plates to be tested the following day, 313 larvae were recovered and retested. The 4 lost larvae were excluded from analysis.

### Cycloheximide feeding protocol

For specified experiments in section **Figure 4**, larva were raised on ATR-food plates at 25C for 48 hours. Second instar larvae were separated from food using 30% sucrose solution and washed in water. Four hours prior to experiments, larvae were transferred to an agar dish with yeast paste for feeding. Yeast paste was made with either a solution of 35 mM cycloheximide (CXM, Sigma Aldrich C7698) and 0.1 mM all-trans-retinal (ATR, Sigma Aldrich R2500) in 5% sucrose (ATR+/CXM+); or 0.1 mM ATR in 5% sucrose (ATR+/CXM-). To verify CXM ingestion, we placed ATR+/CXM+ and ATR+/CXM-larvae not selected for experiments back on clean food and allowed them to continue development. 95% of ATR+/CXM-larvae pupated, while only 45% of ATR+/CXM+ larvae pupated. Before the experiment, larvae were transferred to an empty petri dish and washed with tap water before being placed into a maze. Except where noted, the same experimental protocol was followed as for non-CXM overnight memory.

### Protocol for cycloheximide experiments

For the CXM experiments in section **Figure 4**, larvae were trained with either a massed or spaced training protocol. The 20x massed training protocol was as previously described for other experiments in **Figure 4**.

In the 10x spaced training protocol, larvae were first tested for one hour to determine their initial CO_2_ preference. They then received two cycles of paired DAN-i1 activation with CO_2_ presentation (15 seconds of CO_2_ presentation paired with reward, followed by 15 seconds of air presentation), followed by 15 minutes of air presentation. This was then repeated five times (10 activations total). In these experiments, we did not test the larvae immediately following training but instead removed them to food and tested their preferences the next day only.

The 10x massed training protocol was identical to the 10x spaced training protocol, except training consisted of 10 sequential cycles of paired DAN-i1 activation with CO_2_ presentation (15 seconds of CO_2_ presentation paired with reward, followed by 15 seconds of air presentation, repeated 10 times). As in the spaced training experiments, larvae were removed to food immediately following training, and their preferences were tested the next day only.

### Habituation and extinction protocols

For experiments in **Figure 3**, we used either an extinction or habituation protocol during training. For both types, larvae were tested for one hour to determine their innate CO_2_ preference in the method described above.

For extinction experiments, larvae were trained in the same Y-maze used to measure preference. 30-second training cycles alternate 15 seconds of CO_2_ presentation, where both channels contain a mix of CO_2_ and air; followed by 15 seconds of air presentation, where neither channel had odorized air. Red LEDs were used to activate CsChrimson synchronously with CO_2_ presentation. This training cycle was repeated some number of times (specified for each experiment above). Immediately after training, we presented the larva with 18 cycles of repeated CO_2_/air exposure (15 seconds of CO_2_ followed by 15 seconds of air; repeat) with no reward pairing. After these extinction cycles, larva preference for CO_2_ was tested for one hour.

Habituation experiments were done exactly as for extinction experiments, except that the 18 unrewarded cycles of repeated CO_2_/air exposure were presented immediately prior to the training cycles.

For experiments in **Figure 4**B, we tested each larva’s initial preference for one hour, then presented 3 rewarded paired training cycles. For some larvae (‘Extinction Post-Train’), we then immediately presented 18 extinction cycles, removed the larvae to food overnight as described above, and then tested their preferences for one hour the next day. For another set of larvae (‘Extinction Pre-Test’), we removed the larvae to food immediately following training. The next day, after the larvae were cleaned and inserted into the Y-maze, they were exposed to 18 extinction cycles immediately prior to testing their CO_2_ preferences for one hour.

### Protocol for timing dependence experiments

For experiments in **Figure 1**D, reward presentation was offset from CO_2_ onset. 30-second training cycles alternated 15 seconds of CO_2_ presentation, where both channels contain a mix of CO_2_ and air; followed by 15 seconds of air presentation, where neither channel has odorized air. Red LEDs are used to activate CsChrimson for 15 seconds. For some larvae, reward onset occurred 7.5 seconds after CO_2_ presentation; for others, reward onset occurred 7.5 seconds before CO_2_ presentation. For all experiments of this type, larvae were presented with 20 cycles of training.

For experiments in **Figure 1**E, 75-second training cycles alternated 15 seconds of CO_2_ presentation, where both inlet channels contained a mix of CO_2_ and air with 60 seconds of air presentation. For some larvae, reward presentation occurred immediately following CO_2_ termination for 15 seconds. For others, reward presentation occurred 15 seconds prior to CO_2_ onset, and reward presentation was terminated upon CO_2_ presentation. For a third group of larvae, we rewarded larvae for 15 seconds between two CO_2_ presentations. In this case, 15 seconds of CO_2_ presentation was followed by 15 seconds of reward presentation in the absence of CO_2_, followed by another 15 seconds of CO_2_ presentation. After the second presentation, there was a 30 second air gap before the cycle repeated. For all experiments of these types, larvae were presented with 20 cycles of training.

### Data Analysis

The probability of choosing the CO_2_ containing channel was scored for individual larvae and for populations as

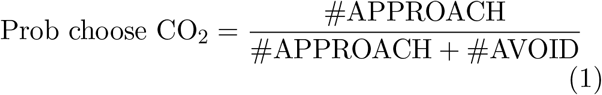

The population average was determined by dividing the total number of times any larva in the population chose the CO_2_ containing channel by the total number of times any larva chose either channel. In other words, larvae that made more decisions contributed more heavily to the average. The number of larva and total number of approach and avoid decisions made by larvae for each type of experiment is shown in **Table S1**.

Error bars and significance tests in the figures were generated by bootstrapping. For each experimental set, we generated 10,000 numerical replicates by selecting with replacement from that set of larvae and then reanalyzing the data. Error bars were the standard deviation of these replicates. A p-value *p < x* indicates that at least *x* fraction of these replicates ended with the same ranking result (e.g. *p <* 0.01 between trained and untrained would indicate that in at least 9900 out of 10,000 replicates, the trained group had a larger CO_2_ preference than the untrained group or vice versa). Note that in each replicate, the same animals are included in each (e.g. trained and untrained) group. In **Table S2**, we also show p-values for the Fisher’s exact test, which treats every decision as independent, and the Mann-Whitney u-test, which treats every larva in each group as a discrete measurement and does not account for differing numbers of decisions made by larvae.

To fit the data in **Figure 2** to various models, we grouped the larvae according to the number of cycles (*n*_*c*_) of training they received. In each group, for each larva we quantified the number of decisions made following training. The number of decisions made by the *j*^*th*^ larva that received *n*_*c*_ cycles of training was *n*(*n*_*c*_, *j*) and the fraction of times the larva chose the CO_2_ containing channel was *p*(*n*_*c*_, *j*).

If all larva chose randomly and independently from the two channels with a fixed probability 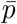 of choosing CO2, then we would expect that the number of times the CO_2_ containing channel would be binomially distributed. For ease of computation, we approximated the binomial distribution as a normal distribution. In this case, the probability density of observing *p*(*n*_*c*_, *j*) given *n*(*n*_*c*_, *j*) would be normally distributed with mean 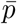 and variance

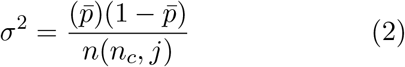

In fact, we found that after choosing a CO_2_ containing channel, both naive and trained larvae are less likely to choose the CO_2_ containing channel the next time they approach the intersection (this effect requires Gr63a so is not an analysis artifact). Because the choices are not independent, the variance of the mean of a series of choices is not given by **Equation 3**. Instead, we modeled the variance as

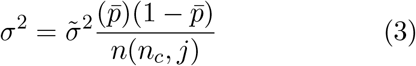

where 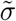 was a global fit parameter in the shifting and exponential fraction models and in the shifting mean model a function of the amount training. This formulation preserved the properties that the variance should increase as the decision to choose CO_2_ approaches 50% and should be larger when fewer decisions are averaged together. However, if we instead just assume a single global *σ*, the results of our analysis (that the exponential fraction model is preferred) are unchanged.

Given this, the probability density of observing an individual larva choosing CO_2_ *p*(*n*_*c*_, *j*) of the time was

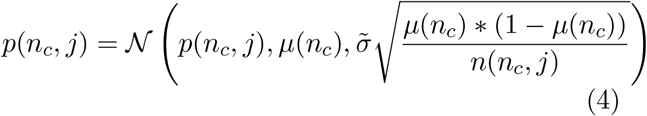

in the shifting mean model, which was represented by a single gaussian whose parameters shifted with the amount of training. In the shifting fraction model,

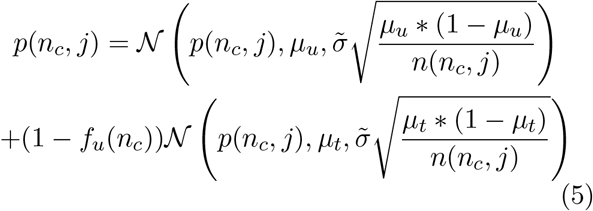

where *f*_*u*_, the fraction of untrained larvae changed with the amount of training, but the means *µ*_*u*_, *µ*_*t*_ and variances of the two populations remained fixed regardless of training.

In the exponential fraction model, the fraction of untrained larvae decreased exponentially.

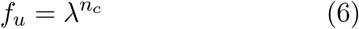

These models were then fit to the data by maximimizing the log-likelihood of the observed data set using the MATLAB function fmincon. The predictions of these fits are shown in **Figure 2**. These results are summarized in **Table S3**, along with the Aikake and Bayes Information Criterion, AIC and BIC, which are used to compare models with different numbers of parameters. According to both AIC and BIC, the exponential fraction model is strongly favored.

## Supplemental Movies

**Figure S1:**
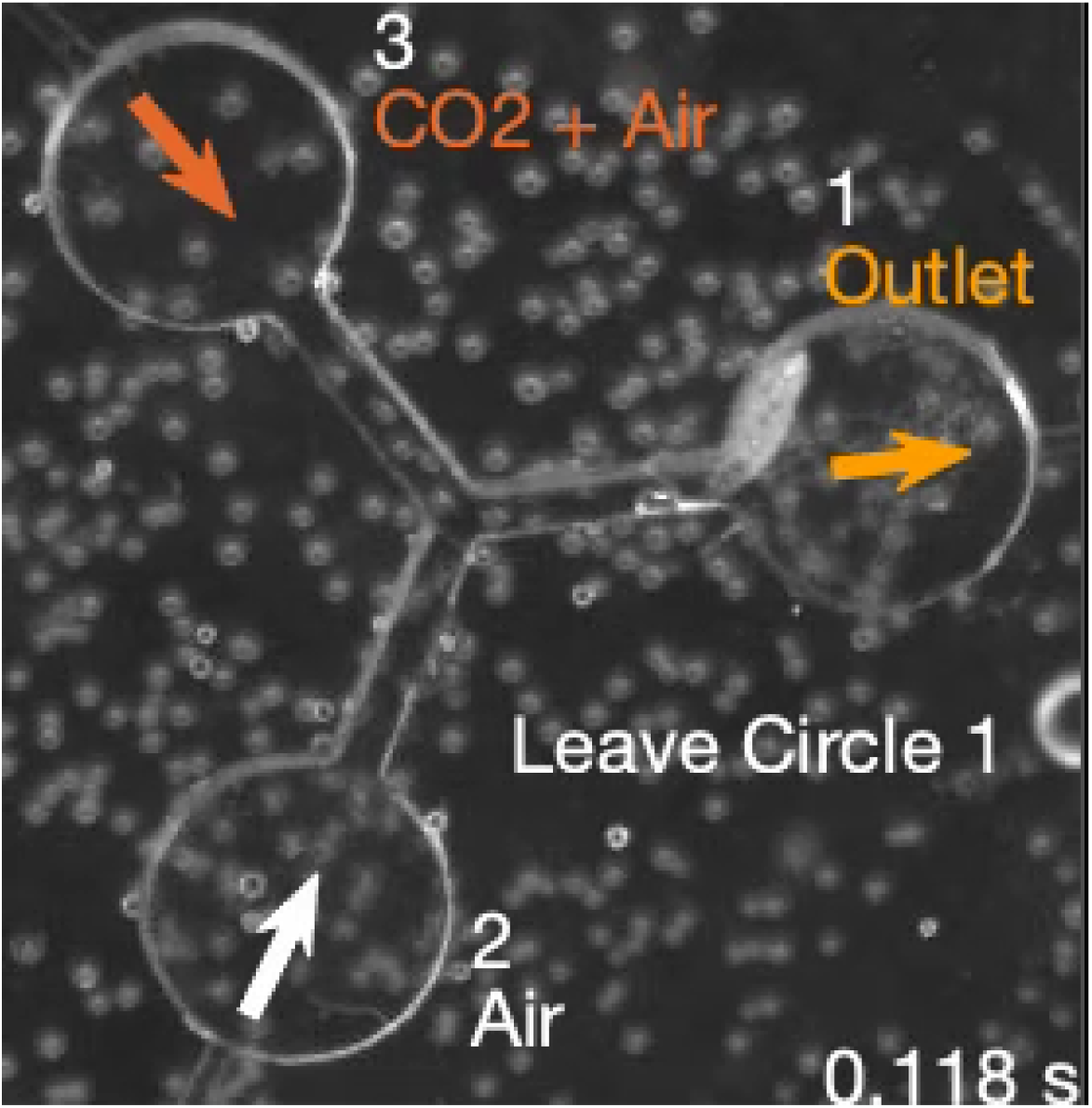
Supplemental Movie 1: Recording of a larva making 2 decisions within the Y-maze. The direction of airflow and the larva’s decisions are noted. Video was recorded at 20 frames per second; the playback speed of 25 fps represents 1.25x real time.

## Supplemental Tables

**Table S1:**
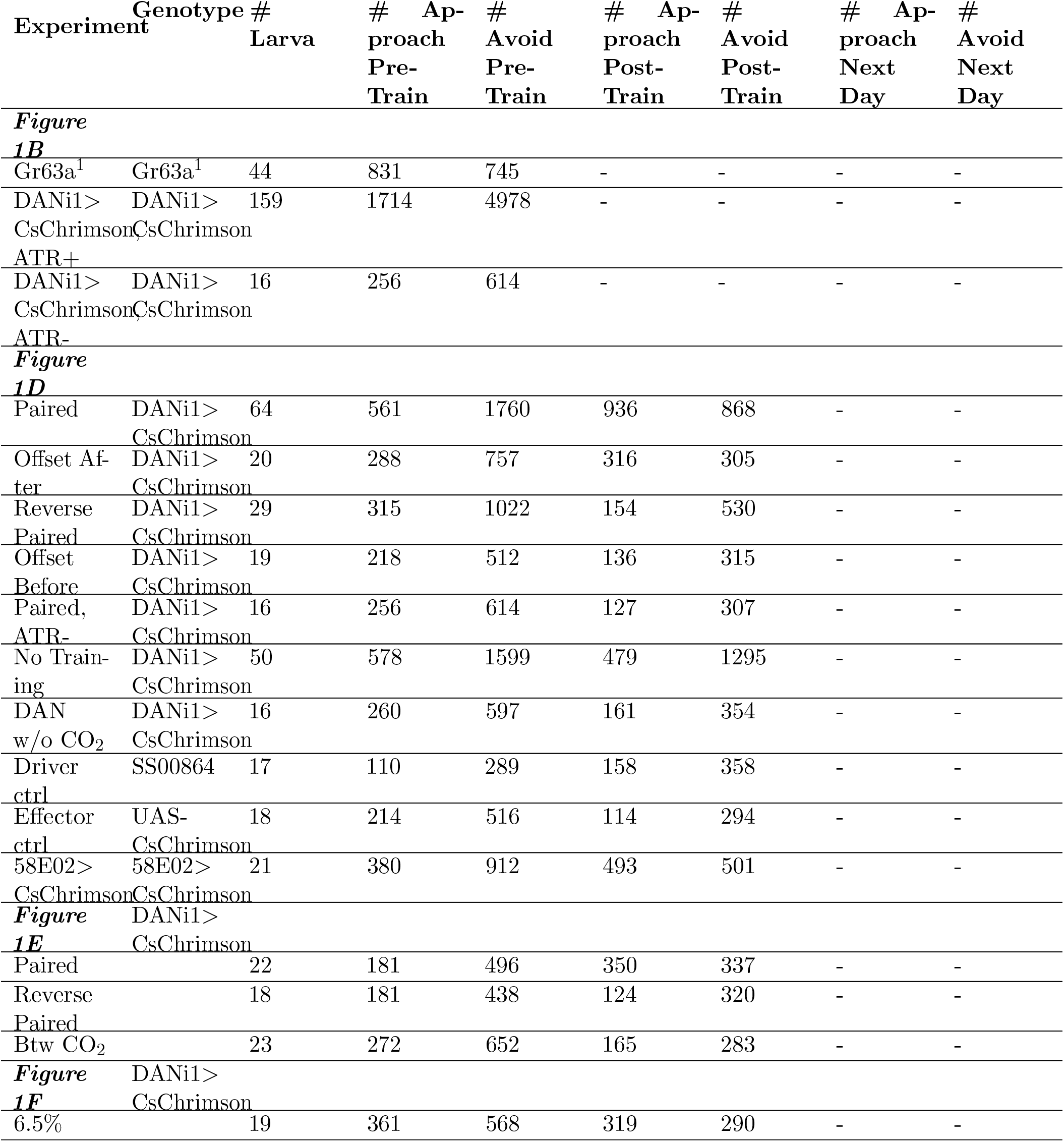
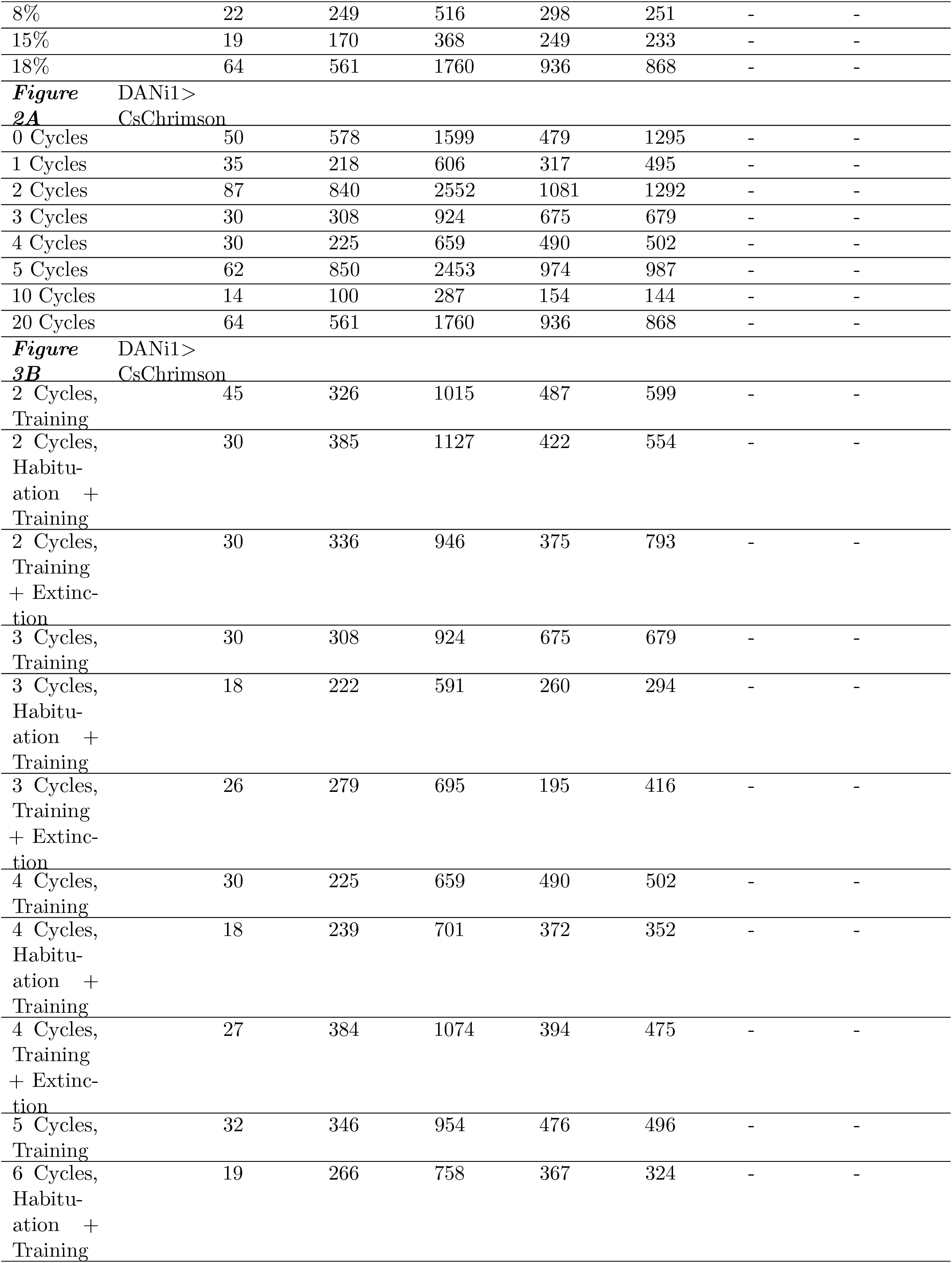
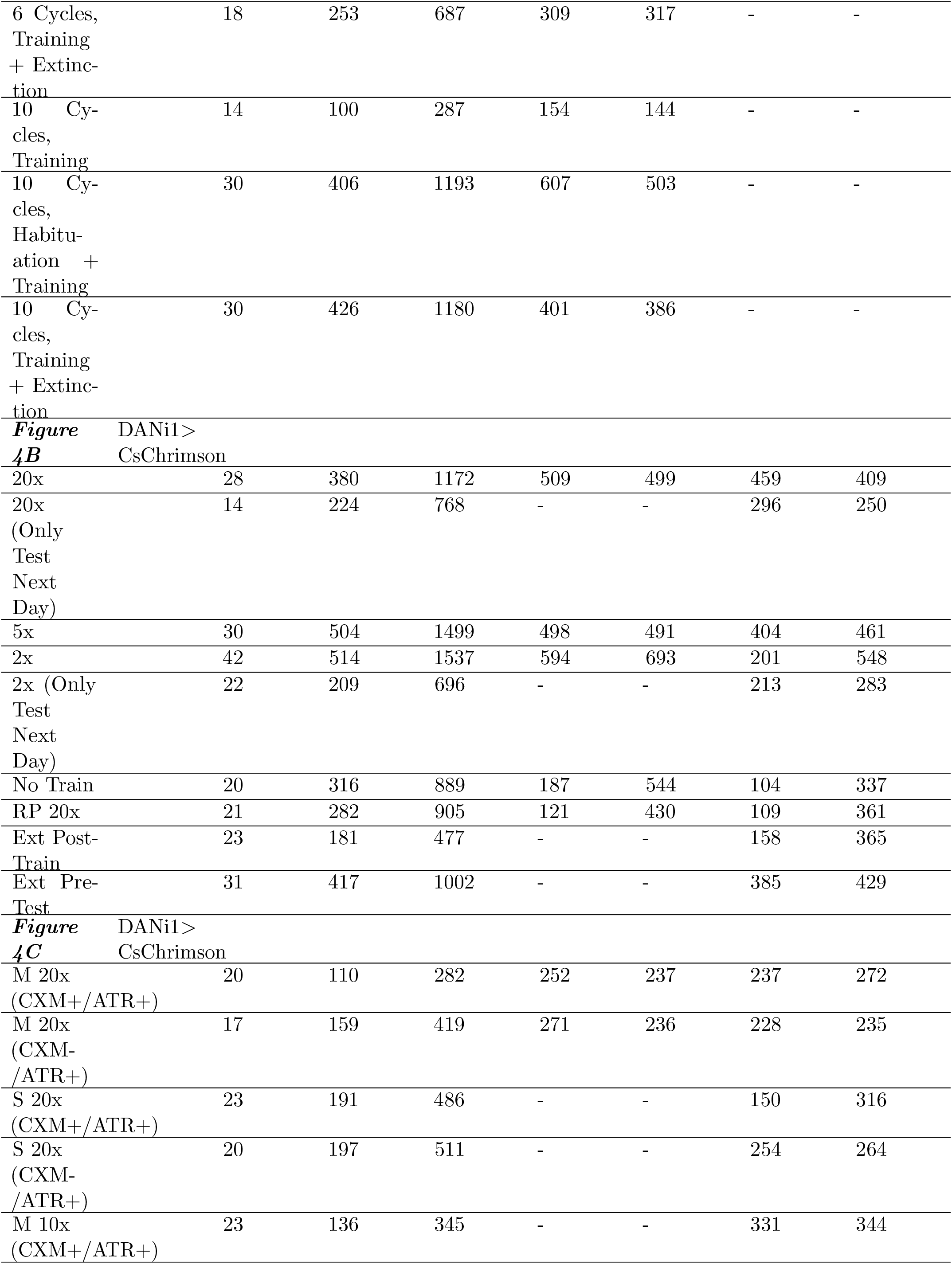
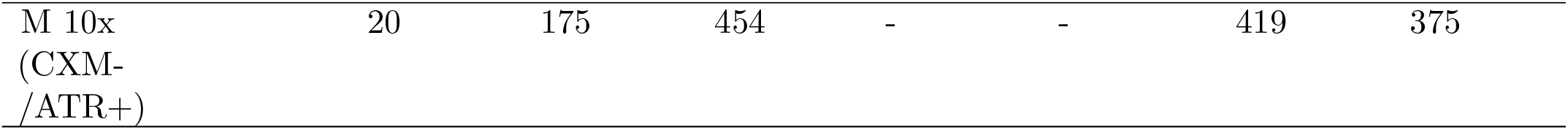
Data for experiments in Figure 1, Figure 2, Figure 3, and Figure 4. # Larva: number of individual larvae tested for experiment type; # Approach Pre-Train: total number of times all larvae chose the channel containing air with CO_2_ prior to training; # Avoid Pre-Train: total number of times all larvae chose the channel containing pure air prior to training; # Approach Post-Train: total number of times all larvae chose the channel containing air with CO_2_ after the indicated training scheme; # Avoid Post-Train: total number of times all larvae chose the channel containing pure air after the indicated training scheme; # Approach Next Day: total number of times all larvae chose the channel containing air with CO_2_ during testing approximately 20 hours after training; # Avoid Next: total number of times all larvae chose the channel containing pure air during testing approximately 20 hours after training. All tests were 1 hour (for each larva).

**Table S2:**
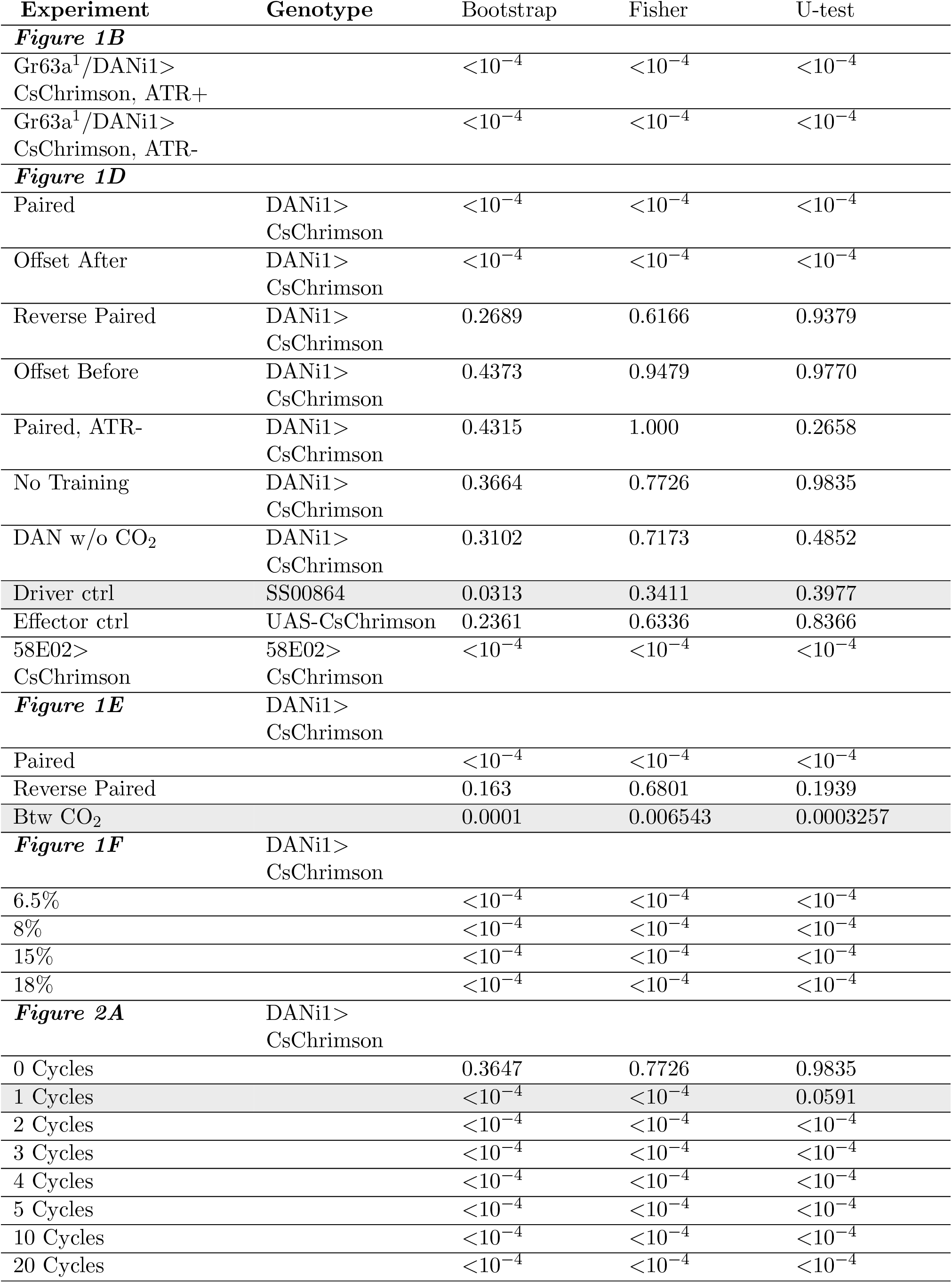
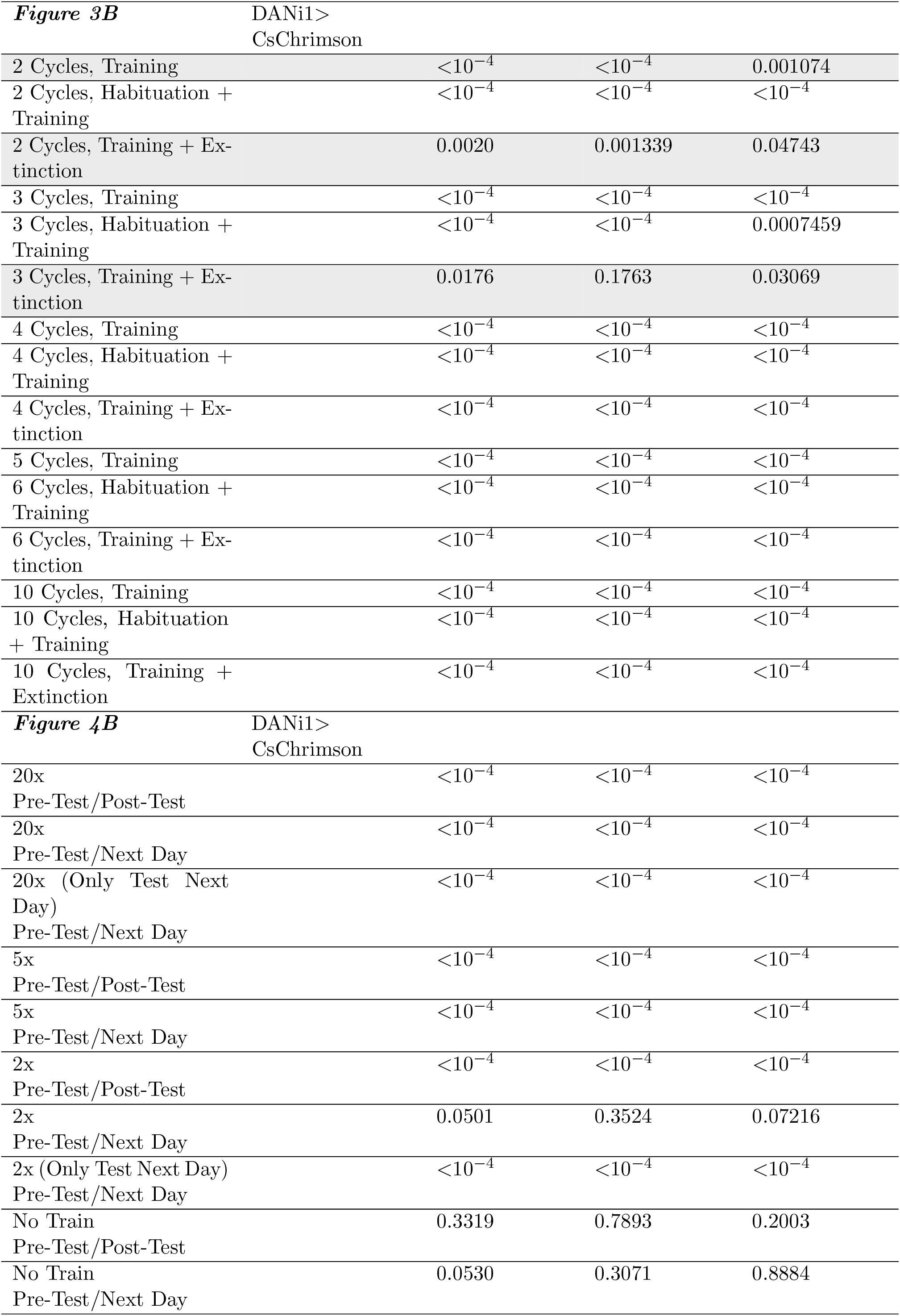
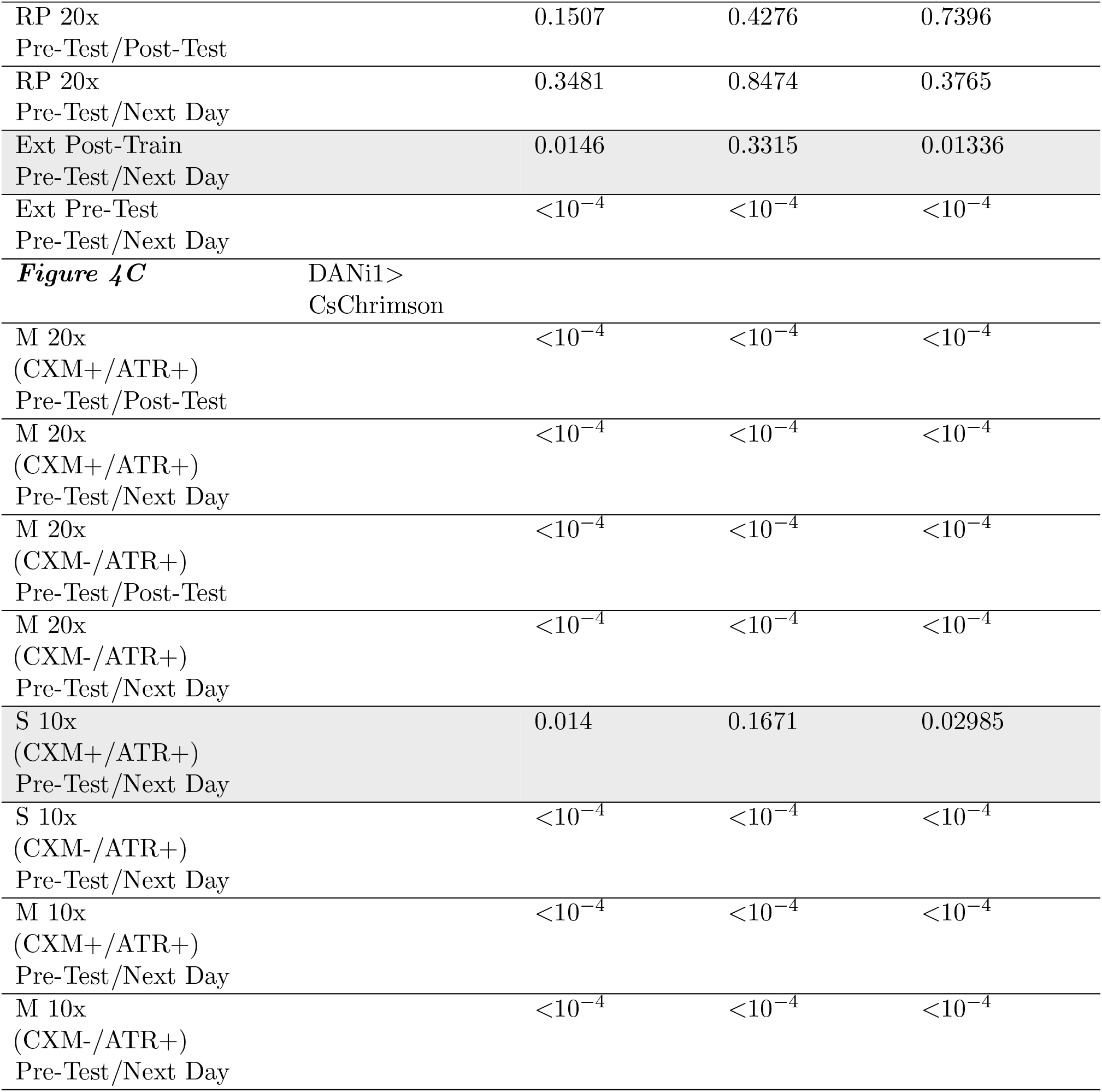
P-values for experiments in Figure 1, Figure 2, Figure 3, and Figure 4. P-values for experiments were calculated: Bootstrap - p-values calculated as explained in Methods; Fisher - p-values calculated using Fisher’s exact test; U-test - p-values calculated using two-sided Mann -Whitney U test. Unless otherwise noted, p-values are calculated between pre-train and post-train data. A shaded row indicates not all tests reach the same significance level (out of ns, p <0.05, p <0.01, p <0.001)

**Table S3:**
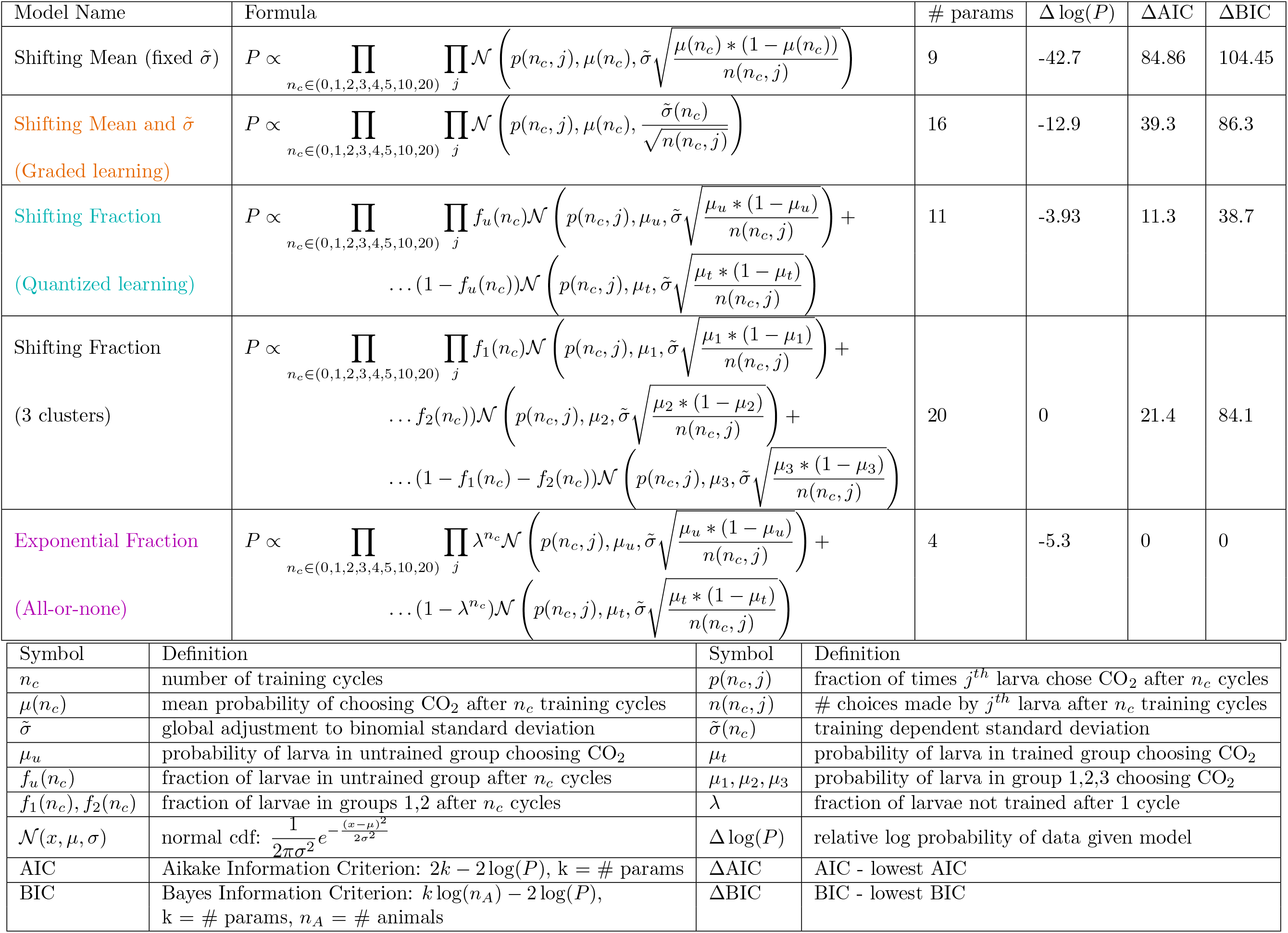
Model fits to data in Figure 2. Shifting Mean and 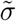, shifting fraction, and exponential fraction models are presented in Figure 2. Model name: name of the model. Formula: expression for the probability of the data given the model and its parameters. # params: number of free parameters in the model. Δ log(P) logarithm of the probability of the data given best fit to this model minus logarithm of the probability of the data given the best fit model overall. ΔAIC, ΔBIC - Aikake and Bayes Information Criterion minus the lowest values over the models tested. Lower numbers indicate model is favored. According to both criterion, the exponential fraction model is strongly favored over the shifting fraction model, and the shifting fraction model is strongly favored over all models except the exponential fractional model.

